# The role of fibroblast growth factor signalling in *Echinococcus multilocularis* development and host-parasite interaction

**DOI:** 10.1101/457168

**Authors:** Sabine Förster, Uriel Koziol, Tina Schäfer, Raphael Duvoisin, Katia Cailliau, Mathieu Vanderstraete, Colette Dissous, Klaus Brehm

## Abstract

**Background:** Alveolar echinococcosis (AE) is a lethal zoonosis caused by the metacestode larva of the tapeworm *Echinococcus multilocularis*. The infection is characterized by tumour-like growth of the metacestode within the host liver, leading to extensive fibrosis and organ-failure. The molecular mechanisms of parasite organ tropism towards the liver and influences of liver cytokines and hormones on parasite development are little studied to date.

**Methodology/Principal findings:** We show that the *E. multilocularis* larval stage expresses three members of the fibroblast growth factor (FGF) receptor family with homology to human FGF receptors. Using the *Xenopus* expression system we demonstrate that all three *Echinococcus* FGF receptors are activated in response to human acidic and basic FGF, which are present in the liver. In all three cases, activation could be prevented by addition of the tyrosine kinase inhibitor BIBF 1120, which is used to treat human cancer. At physiological concentrations, acidic and basic FGF significantly stimulated the formation of metacestode vesicles from parasite stem cells *in vitro* and supported metacestode growth. Furthermore, the parasite’s mitogen activated protein kinase signalling system was stimulated upon addition of human FGF. The survival of metacestode vesicles and parasite stem cells were drastically affected *in vitro* in the presence of BIBF 1120.

**Conclusions/Significance:** Our data indicate that mammalian FGF, which is present in the liver and upregulated during fibrosis, supports the establishment of the *Echinococcus* metacestode during AE by acting on an evolutionarily conserved parasite FGF signalling system. These data are valuable for understanding molecular mechanisms of organ tropism and host-parasite interaction in AE. Furthermore, our data indicate that the parasite’s FGF signalling systems are promising targets for the development of novel drugs against AE.

**Author summary:** To ensure proper communication between their different cell populations, animals rely on secreted hormones and cytokines that act on receptors of target cells. Most of the respective cytokines, such as FGFs, evolved over 500 million years ago and are present in similar form in all animals, including parasitic worms. The authors of this study show that the metacestode larva of the tapeworm *E. multilocularis*, which grows like a malignant tumor within the host liver, expresses molecules with homology to FGF receptors from mammals. The authors show that human FGF, which is abundantly present in the liver, stimulates metacestode development and that all parasite FGF receptors are activated by human FGF, despite 500 million years of evolutionary distance between both systems. This indicates that cells of the *Echinococcus* metacestode can directly communicate with cells of the mammalian host using evolutionarily conserved signaling molecules. This mode of host-pathogen interaction is unique for helminths and does not occur between mammals and single-celled pathogens such as protozoans or bacteria. The authors finally demonstrate that BIBF 1120, a drug used to treat human cancer, targets the *Echinococcus* FGF receptors and leads to parasite death. This opens new ways for the development of anti-parasitic drugs.

## Introduction

The flatworm parasite *E. multilocularis* (fox-tapeworm) is the causative agent of alveolar echinococcosis (AE), one of the most dangerous zoonoses of the Northern hemisphere. Infections of intermediate hosts (rodents, humans) are initiated by oral uptake of infectious eggs which contain the parasite’s oncosphere larval stage [1,2]. After hatching in the host intestine and penetration of the intestinal wall the oncosphere gains access to the liver, where it undergoes a metamorphotic transition towards the metacestode larval stage [3]. The *E. multilocularis* metacestode consists of numerous vesicles which grow infiltratively, like a malignant tumor, into the liver tissue [1-3]. Due to the unrestricted growth of the metacestode, blood vessels and bile ducts of the liver tissue of the intermediate host are obstructed, eventually leading to organ failure [1]. Another hallmark of AE is extensive liver fibrosis which can lead to a complete disappearance of the liver parenchyma, and which most probably involves the activation of hepatic stellate cells during chronic infection [4,5]. Surgical removal of the parasite tissue, the only possible cure, is not feasible in the majority of patients leaving benzimidazole-based chemotherapy as the only treatment option. However, benzimidazoles act parasitostatic only and have to be given for prolonged periods of time (often life-long), underscoring the need for novel treatment options against AE [1].

We previously established that *E. multilocularis* development and larval growth is exclusively driven by a population of somatic stem cells, the germinative cells, which are the only mitotically active cells of the parasite and which give rise to all differentiated cells [6]. Using *in vitro* cultivation systems for metacestode vesicles and germinative cells [7-10], we also demonstrated that host insulin fosters parasite development by acting on evolutionarily conserved receptor kinases of the insulin receptor family that are expressed by the metacestode [11]. Evidence has also been obtained that host epidermal growth factor (EGF) stimulates *Echinococcus* germinative cell proliferation, most probably by acting on parasite receptor tyrosine kinases of the EGF receptor family [12,13]. These studies indicate that the interaction of host-derived hormones and cytokines with corresponding receptors of evolutionarily conserved signalling pathways that are expressed by the parasite may play an important role in AE host-parasite interaction. Although the *E. multilocularis* genome project already indicated that the parasite expresses receptor tyrosine kinases of the fibroblast growth factor (FGF) receptor family in addition to insulin- and EGF-receptors [14], no studies concerning the effects of host FGF and their possible interaction with parasite FGF receptors have been carried out to date.

FGFs are an ancient group of polypeptide cytokines that are present in diploblastic animals, in deuterostomes and, among protostomes, only in ecdysozoa (with some distantly related members in lophotrochozoa)[15,16]. Humans express 22 different FGFs of which several (FGF11 – FGF14) are not secreted and act independently of FGF receptors in an intracrine modus only [15]. The remaining FGFs act in a paracrine fashion and are typically released via N-terminal signal peptides. Notable exceptions are the prototypic FGF1 (acidic FGF) and FGF2 (basic FGF) which are ubiquitously expressed in human tissues, are the most active members of the FGF family, and are released in a signal peptide-independent manner [15]. FGFs have a key role in metazoan embryonic development and, in adults, are typically involved in regeneration processes (angiogenesis, wound healing, liver regeneration, regeneration of nervous tissue)[15]. In the liver, particularly FGF1 but also FGF2 are present as proteins in significant amounts [17], are crucially involved in tissue regeneration upon damage [18,19], and are also upregulated and released during fibrosis [20]. Secreted FGFs act through surface receptor tyrosine kinases of the FGF receptor family, of which four isoforms, Fgfr1-Fgfr4, are expressed by humans [15]. The mammalian FGF receptors comprise an extracellular ligand-binding domain made up of three immunoglobulin (Ig)-like domains, a transmembrane domain, and a split intracellular kinase domain. FGF binding to the cognate FGF receptors typically results in receptor dimerization, transphosphorylation and subsequent activation of downstream signalling pathways such as the Ras-Raf-MAPK (mitogen-activated protein kinase) cascade or the PI3K/Akt pathway [15].

FGF signalling pathways have, in part, already been studied in flatworms. In free-living planarian species *Dugesia japonica*, two members of the FGFR tyrosine kinases are expressed of which DjFGFR1 exhibits three immunoglobulin-like domains in the extracellular region whereas DjFGFR2 only contains two such domains [21]. Both receptors are expressed by X-ray sensitive planarian stem cells (neoblasts) and in cephalic ganglia and an important role of these FGFRs in planarian brain formation has been suggested [21,22]. Furthermore, similar FGF receptors were also detected in stem cells of the planarian *Schmidtea mediterranea* [23]. In the genome of the flatworm parasite species *Schistosoma mansoni*, two FGFR-encoding genes were identified of which *fgfr*A codes for a predicted protein with two extracellular immunoglobulin domains and a split tyrosine kinase domain whereas the *fgfr*B gene product only comprises one immunoglobulin domain in the extracellular region [24]. Expression of *fgfr*A and *fgfr*B in neoblast-like somatic stem cells has been shown and evidence was obtained for an important role of both receptors in schistosome stem cell maintenance [25-27]. Hahnel et al. [24] also demonstrated that both receptors are enzymatically active, are expressed in the gonads of schistosomes, and are upregulated following pairing, indicating a role in parasite fertility. Interestingly, these authors also showed that treatment of adult schistosomes with FGFR inhibitors leads to a reduction of somatic neoblast-like stem cells in both genders [24].

In the present work we provide a detailed analysis of three FGFRs in the cestode *E. multilocularis* and show that the expression patterns of these receptors differ from those in planaria and schistosomes. We also demonstrate that all three *Echinococcus* FGFRs are activated in response to human FGFs and that host FGF stimulates parasite development *in vitro*. Finally, we also show that inhibition of FGF signalling in *Echinococcus* larvae drastically reduces parasite development and survival.

## Methods

### Organisms and culture methods

FGF stimulation and inhibitor experiments were performed with the natural *E. multilocularis* isolate H95 [14]. Whole mount in situ hybridization was carried out using isolate GH09 which, in contrast to H95, is still capable of producing brood capsules and protoscoleces *in vitro* [14]. All isolates were continuously passaged in mongolian jirds (*Meriones unguiculatus*) as previously described [9]. The generation of metacestode vesicles and axenic cultivation of mature vesicles was performed essentially as previously described [7,9] with media changes usually every three days. Primary cell cultures were isolated from mature vesicles of isolate H95 and propagated *in vitro* essentially as previously described [8-10] with media changes every three days unless indicated otherwise. For FGF stimulation assays, 10 nM or 100 nM of recombinant human acidic FGF (FGF1) or basic FGF (FGF2)(both from ImmunoTools GmbH, Friesoythe, Germany) were freshly added to parasite cultures during medium changes. In the case of primary cells, cultivation was usually performed in cMEM medium which is host hepatocyte-conditioned DMEM (prepared as described in [10]). For inhibitor studies, specific concentrations of BIBF 1120 (Selleck Chemicals LLC, Houston, TX, USA) were added to parasite cultures as indicated and as negative control DMSO (0.1 %) was used. The formation of mature metacestode vesicles from primary cells and measurement of metacestode vesicles size was performed essentially as previously described [11].

### Nucleic acid isolation, cloning and sequencing

RNA isolation from in vitro cultivated axenic metacestode vesicles (isolate H95), protoscoleces (isolate GH09), and primary cells (H95, GH09) was performed using a Trizol (5Prime, Hamburg, Germany)-based method as previously described [11]. For reverse transcription, 2 µg total RNA was used and cDNA synthesis was performed using oligonucleotide CD3-RT (5’-ATC TCT TGA AAG GAT CCT GCA GGT_26_ V-3’). PCR products were cloned using the PCR cloning Kit (QIAGEN, Hilden, Germany) or the TOPO XL cloning Kit (invitrogen), and sequenced employing an ABI prism 377 DNA sequencer (Perkin-Elmer).

The full-length *emfr1* cDNA was cloned using as starting material the partial sequence of a cDNA of the closely related cestode *E. granulosus*, which encoded a FGFR-like tyrosine kinase domain but which lacked the coding regions for transmembrane and extracellular parts [28]. Using primers directed against the *E. granulosus* sequence (5’-CTA CGC GTG CGT TTT CTG ATG-3’for first PCR; 5’-CCC TCT GAT CCA ACC TAC GAG-3’for nested PCR), the 3’ end of the corresponding *E. multilocularis* cDNA was subsequently PCR amplified from a metacestode (isolate H95) cDNA preparation using primers CD3 and CD3nest as previously described [29]. 5’-RACE was performed using the SMART RACE cDNA amplification kit (Clontech) according to the manufacturer’s instructions using primers 5’-ACC GTA TTT GGG TTG TGG TCG-3’ (first PCR) and 5’-GAA CAG GCA GAT CGG CAG-3’ (touchdown PCR) as previously described [30]. The presence of an in frame TAA stop codon 110 bp upstream of the *emfr1* ATG start codon indicated that the correct 5’ end had been identified. In a final step, the entire *emfr1* cDNA was PCR amplified from metacestode cDNA using primers 5’-GAC ACA TCT CCT TGG CCG-3’ and 5’-GCG AGT TGA TAC TTT ATG AGA G-3’ and cloned using the TOPO XL PCR cloning kit (Invitrogen). The sequence is available in the GenBank™, EMBL, and DDJB databases under the accession number LT599044.

For *emfr2* cloning we first identified by BLAST analyses on the published *E. multilocularis* genome sequence [14] a reading frame encoding a FGFR-like TKD annotated as EmuJ_000196200. Transcriptome analyses [14] and 5’_RACE experiments, however, indicated that there is actually read-through transcription between gene models EmuJ_000196300 and EmuJ_000196200. We thus designed primers 5’-ATG TGT CTC CGA GCT CTC TG-3’, binding to the 5’ end regions of gene model EmuJ_000196300, and primer 5’-TTA CTC GCT CGA TCG TGG GG-3’, binding to the reading frame 3’ end of gene model EmuJ_000196200, to PCR amplify the entire reading frame from metacestode cDNA. The resulting PCR fragment was subsequently cloned using the TOPO XL cloning kit (Invitrogen) and fully sequenced. The sequence is available in the GenBank™, EMBL, and DDJB databases under the accession number LT599045.

For *emfr3* cloning and sequencing we used primers directed against the CDS 5’ end (5’-ATG GCA CCT AAG GTT GTG TCA GGA-3’) and 3’ end (5’-GCA GAT GAG TAA GAA ACC CTC-3’) of gene model EmuJ_000893600 [14] for direct PCR amplification of the reading frame from metacestode cDNA. The resulting PCR fragment was subsequently cloned using the TOPO XL cloning kit (Invitrogen) and sequenced. The sequence is available in the GenBank™, EMBL, and DDJB databases under the accession number LT599046.

### BrdU proliferation assays

Proliferation of *E. multilocularis* metacestode vesicles and primary cells was assessed by a bromodesoxyuridine (BrdU)-based method. Axenically cultivated metacestode vesicles (2-4 mm in diameter) were manually picked and incubated in 12-well plates (Greiner BioOne, Kremsmünster, Germany; 8 vesicles per well) in DMEM medium without serum for 2 days. Freshly isolated primary cells were plated on 12-well plates and grown for 2 days under axenic conditions in conditioned DMEM (cMEM) medium with serum [8]. BrdU (SigmaAldrich, taufkirchen, Germany) as well as recombinant human FGF1 and FGF2 were added at 1mM (BrdU) and 100 nM or 10 nM (FGF1, FGF2) final concentrations as indicated. Cultures were incubated for 48 h at 37°C under 5% CO_2_ for metacestode vesicles or under nitrogen atmosphere [7,8] in the case of primary cells. Samples were analysed in duplicates in three independent experiments. As controls, metacestode vesicles or primary cells were incubated in either DMEM without serum or conditioned DMEM, without addition of FGFs.

Primary cells and metacestode vesicles were then isolated for genomic DNA analysis. In detail, vesicles and primary cells were first washed with 1xPBS, pelleted, and subsequently transferred to lysis buffer (100 mM NaCl, 10 mM Tris-HCl, pH 8.0; 50 mM EDTA, pH 8.0, 0,5% SDS) supplemented with 20 μg/ml RNAse A and 0,1 mg/ml proteinase K. Samples were then incubated at 50°C for 4 h under constant shaking for complete lysis. DNA was isolated by two rounds of phenol/chlorophorm extraction (1 vol of phenol/chlorophorm/isoamylalcohol 25:24:1). DNA was then precipitated with 2 vol of 96% ethanol and 0,1 vol of LiCl (pH 4,5) after overnight incubation at −20°C and centrifugation at 20.000 rcf for 30 min at 4°C and washed with 70% ethanol. The pellet was then air dried for 15 min an resuspended in 1 x TE buffer (10 mM Tris, 1 mM EDTA, pH 8,0).

The DNA was then prepared for coating onto a 96-well plate (96 well optical bottom plates, Nunc, Langenselbold, Germany). To this end, 5 µg of DNA were combined with 1 vol of Reacti-Bind DNA Coating solution (Pierce Biotechnology, Rockford, IL, USA) and mixed for 10 min. The DNA mix was then added to the microplates in duplicates and incubated overnight at room temperature with gentle agitation. The TE/Reacti-Bind DNA coating solution mix served as a negative control. Unbound DNA was removed by washing three times with 1xPBS. After blocking with 5% nonfat dry milk in 1xPBS for 1 h at room temperature and extensive washing with 1xPBS, 100 µl of anti-BrdU-POD (Cell Proliferation ELISA, BrdU; Roche Applied Science, Mannheim, Germany) was added and incubated for 90 min at room temperature. After incubation, microplates were washed three times with 1xPBS buffer before substrate solution (Cell Proliferation ELISA, BrdU; Roche Applied Science, Mannheim, Germany) was added and the wells were incubated for 60 min. Stop-solution (25 µl of 1 M H_2_SO_4_) was added and absorbance of the samples was measured using an ELISA reader at 450 nm.

### In situ hybridization protocols

Coding sequences of FGF receptors from *E. multilocularis* were amplified by RT-PCR and cloned into the vectors pDrive (Qiagen) or pJET1.2 (Thermo Fisher). In the case of *emfr2*, the full length coding sequence was amplified using primers 5’-ATG TGT CTC CGA GCT CTC TG-3’(forward) and 5’-TTA CTC GCT CGA TCG TGG GG-3’ (reverse), whereas partial coding sequences were amplified for *emfr1* (using forward primer 5’-GCA GTG GGC GTC TTC TTT CAC-3’ and reverse primer 5’-GTA AAT GTG GGC CGA CAC TCA G-3’) and for *emfr3* (using forward primer 5’-TTG CCC AGT CAT CCG CTA CAA G-3’ and reverse primer 5’-GCA AGC GGT CAT GAG GCT GTA G-3’). The recombinant plasmids were used for *in vitro* transcription of digoxigenin-labelled RNA probes as previously described [6]. These probes were used for fluorescent WMISH of in vitro cultured *E. multilocularis* larvae as described in [6]. Control WMISH experiments using the corresponding sense probes were always negative (see Figure 2 and data not shown).

### *Xenopus* oocyte expression assays

For expression in the *Xenopus* system, the *emfr1*, *emfr2*, and *emfr3* coding sequences without predicted signal peptide information were cloned into the pSecTag2/Hygro expression system (ThermoFisher Scientific, Germany) leading to an in frame fusion of the Igk leader sequence (provided by the vector system) and the *E. multilocularis* FGF receptor sequences under control of the T7 promoter. Capped messenger RNAs (cRNA) encoding EmFR1, EmFR2 and EmFR3 were then synthesized *in vitro* using the T7 mMessage mMachine Kit (Ambion, USA). Microinjection of EmFGFR cRNAs (60 ng in 60 µl) was performed in stage VI *Xenopus laevis* oocytes according to the procedure previously described [31]. Following 48h of receptor expression, human FGF1 or FGF2 (R & D systems, UK) were added to the extracellular medium at the final concentration of 10 nM. cRNA of *Pleurodeles* FGFR1 identified as homologous to human receptor [32] was a gift of Shi D.L. (CNRS UMR 722, Paris VI) and was used as a positive control.

In some experiments, BIBF1120 (stock solution 10mM in DMSO, Selleck Chemicals LLC) was added (0.1 to 20 µM final concentration) 1 h before the addition of 10 nM FGF1 or FGF2 on EmFR1, EmFR2, EmFR3 and *Pleurodeles* FGFR1 expressing oocytes. Following 15 h of FGF1 or FGF2 stimulation, oocytes were analyzed for their state of progression in the cell cycle. The detection of a white spot at the animal pole of the oocyte attested to G2/M transition and GVBD. Non-injected oocytes treated with progesterone (10 µM) were used as positive controls of GVBD. For each assay, sets of 20-30 oocytes removed from 3 different animals were used.

Dead kinase (TK-) receptors were obtained by site-directed mutagenesis of the EmFR1, EmFR2 and EmFR3 constructs. The active DFG sites present in EmFR1 (D_442_FG), EmFR2 (D_647_FG) and EmFR3 (D_701_FG) were replaced by an inactivating motif (DNA) as described in [31].

For western blot analysis, oocytes were lysed in buffer A (50 mM Hepes pH 7.4, 500 mM NaCl, 0.05% SDS, 5 mM MgCl2, 1 mg ml−1 bovine serum albumin, 10 μg ml−1 leupeptin, 10 μg ml−1 aprotinin, 10 μg ml−1 soybean trypsin inhibitor, 10 μg ml−1 benzamidine, 1 mM PMSF, 1 mM sodium vanadate) and centrifuged at 4 °C for 15 min at 10,000 g. Membrane pellets were resuspended and incubated for 15 min at 4 °C in buffer A supplemented with 1% Triton X-100 and then centrifuged under the same conditions. Supernatants were analyzed by SDS-PAGE. Proteins were transferred to a Hybond ECL membrane (Amersham Biosciences, France). Membranes were incubated with anti-myc (1/50 000, Invitrogen France) or anti-PTyr (1/8000, BD Biosciences, France) antibodies and secondary anti-mouse antibodies (1/50 000, Biorad, France). Signals were detected by the ECL advance Western blotting detection kit (Amersham Biosciences, France)

### MAPK cascade activation assays

Axenically cultivated metacestode vesicles of about 0.5 cm in diameter were incubated in DMEM medium with or without 10% FCS for 4 days. Vesicles cultivated without FCS were subsequently incubated with 10 nM FGF1 (aFGF) or 10 nM FGF2 (bFGF) for 30 sec, 60 sec or 60 min. Immediately after stimulation, vesicles were harvested, cut by a scalpel to remove cyst fluid and then subjected to protein isolation as described previously [33]. Isolated protein lysates were then separated on a 12% acrylamide gel and analysed by Western blotting using a polyclonal anti-Erk1/Erk2 antibody (ThermoFisher Scientific; #61-7400), recognizing Erk-like MAP kinases in phosphorylated and non-phosphorylated form, as well as a polyclonal antibody against phospho-Erk1/Erk2 (ThermoFisher Scientific; #44-680G), specifically directed against the double-phosphorylated (activated) form of Erk1/Erk2 (Thr-185, Tyr-187). We had previously shown that these antibodies also recognize the Erk-like MAP kinase EmMPK1 of *E. multilocularis* in phosphorylated and non-phosphorylated form [33]. As secondary antibody, a peroxidase-conjugated anti-mouse IgG antibody (Dianova, Hamburg, Germany) was used. In inhibitor experiments, axenically cultivated metacestode vesicles were incubated with either 5 µM or 10 µM BIBF 1120 for 30 min and then processed essentially as described above.

### Computer analyses and statistics

Amino acid comparisons were performed using BLAST on the nr-aa and swissprot database collections available under (https://www.genome.jp/). Genomic analyses and BLAST searches against the *E. multilocularis* genome [14] were done using resources at (https://parasite.wormbase.org/index.html). CLUSTAL W alignments were generated using MegAlign software (DNASTAR Version 12.0.0) applying the BLOSUM62 matrix. Domain predictions were carried out using the simple modular architecture research tool (SMART) available under (http://smart.embl-heidelberg.de/) as well as PROSITE scans available under (https://prosite.expasy.org/scanprosite/). Two-tailed, unpaired student’s T-tests were performed for statistical analyses (GraphPad Prism, version 4). Error bars represent standard error of the mean. Differences were considered significant for p-values below 0.05 (indicated by *).

### Ethical approval

All experiments were carried out in accordance with European and German regulations on the protection of animals (*Tierschutzgesetz*). Ethical approval of the study was obtained from the local ethics committee of the government of Lower Franconia (permit no. 55.2 DMS 2532-2-354).

## Results

### The *E. multilocularis* genome encodes FGF receptor kinases but no canonical FGF ligands

By cDNA library screening and mining of the available *E. multilocularis* genome sequence we identified a total of three *Echinococcus* genes encoding members of the FGFR family of receptor tyrosine kinases. A partial cDNA for a gene encoding a tyrosine kinase with homology to FGFRs was previously cloned for *E. granulosus* [28] and by RT-PCR amplification of metacestode cDNA as well as 5’-RACE, the entire cDNA of the *E. multilocularis* ortholog, designated *emfr1* (*<u>E</u>. <u>m</u>ultilocularis* <u>f</u>ibroblast growth factor <u>r</u>eceptor <u>1</u>), was subsequently cloned by us in this work. As shown in Fig 1, the encoded protein, EmFR1, contained an N-terminal export directing signal peptide, followed by one single Ig-like domain, a transmembrane region, and an intracellular tyrosine kinase domain (Figs 1 and S1). In the recently released *E. multilocularis* genome information [14], this gene was correctly predicted on the basis of genome and transcriptome data (*E. multilocularis* gene designation: EmuJ_000833200). In the upstream genomic regions of *emfr1*, no information encoding potential Ig-like domains was identified which, together with the presence of a signal peptide sequence upstream of the single Ig-like domain, indicated that EmFR1 indeed contained only one single Ig-like domain. Amino acid sequence alignments indicated that the kinase domain of EmFR1 contains all residues critical for enzymatic activity at the corresponding positions (Fig S2) and SWISS-PROT database searches revealed highest similarity between the EmFR1 kinase domain and that of human FGFR4 (42% identical aa; 59% similar aa).

**Fig 1.**
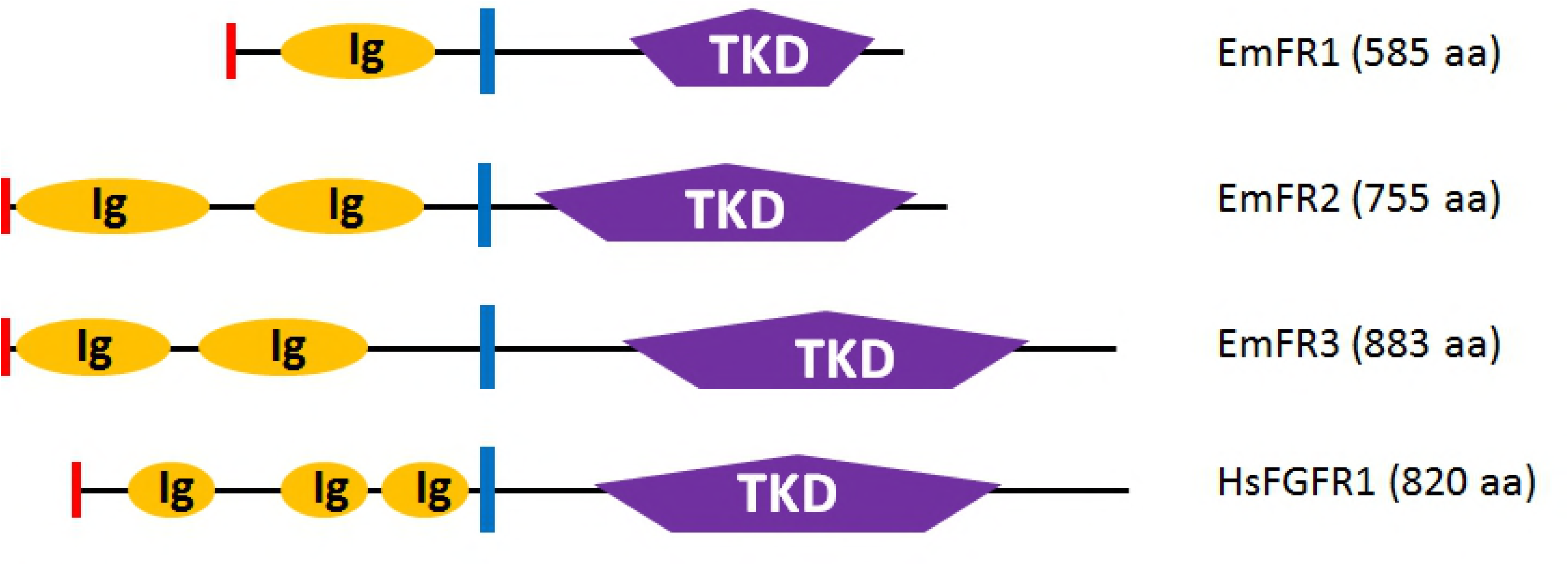
Domain structure of *E. multilocularis* FGF receptors. Schematic representation of the domain structure of the *E. multilocularis* receptors EmFR1, EmFR2, and EmFR3 according to SMART (Simple Modular Architecture Tool) analyses. As a comparison, the structure of the human FGFR1 (HsFGFR1) is shown. Displayed are the location and size of the following predicted domains: tyrosine kinase domain (TKD) in purple and IG-domain (Ig) in orange. Predicted signal peptides are depicted in red and transmembrane domains are shown as blue bars. The numbers of amino acids in full length receptors are shown to the right.

A second gene encoding a tyrosine kinase with significant homology to known FGFRs was identified on the available *E. multilocularis* genome sequence [14] under the annotation EmuJ_000196200. The amino acid sequence of the predicted protein only contained an intracellular tyrosine kinase domain, a transmembrane region, and one extracellular Ig-like domain, but no putative signal peptide. We therefore carried out 5’-RACE analyses on a cDNA preparation deriving from protoscolex RNA and identified the remaining 5’ portion of the cDNA, which contained one additional Ig-like domain and a predicted signal peptide. In the genome annotation, these remaining parts were wrongly annotated as a separate gene under the designation EmuJ_000196300. Hence, the second FGFR encoding gene of *E. multilocularis*, *emfr2*, encoding the protein EmFR2, actually comprises the gene models EmuJ_000196300 and EmuJ_000196200 of the genome sequence. EmFR2 thus comprises a signal peptide, two extracellular Ig-like domains, a transmembrane region, and an intracellular TKD (Figs 1 and S1). The TKD contained all residues critical for tyrosine kinase activity (Fig S2) and, in SWISS-PROT BLASTP analyses, showed highest similarity to two FGF receptor kinases of the flatworm *Dugesia japonica* (45% identical, 65% similar residues), to the *S. mansoni* receptor FGFRB (55%, 68%) and to human FGFR3 (48%, 62%).

The third FGF receptor encoding gene of *E. multilocularis*, *emfr3*, was identified under the designation EmuJ_000893600 and was originally listed as an ortholog of the tyrosine protein kinase Fes:Fps [14]. However, unlike the Fes:Fps kinase which contains FCH and SH2 domains, the EmuJ_000893600 gene product, EmFR3, comprised an N-terminal signal peptide, two extracellular Ig-like domains, a transmembrane region, and an intracellular TKD (Figs 1 and S1), in which 22 of 30 highly conserved residues of tyrosine kinases are present at the corresponding position (Fig S2). Furthermore in SWISS-PROT BLASTP analyses the EmFR3 TKD displayed highest similarity to several vertebrate FGF receptors and to human FGFR2 (32%, 47%). We thus concluded that EmuJ_000893600 actually encoded a third *Echinococcus* FGF receptor tyrosine kinase.

Apart from *emfr1*, *emfr2*, and *emfr3*, no further genes were identified in the *E. multilocularis* genome which displayed clear homology to known FGFR TKDs and which contained characteristic IG domains in the extracellular portions. In vertebrates, structural homology has been described between the TKDs of the receptor families of FGF receptors, the vascular endothelial growth factor (VEGF) receptors, and the platelet-derived growth factor (PDGF) receptors, which also contain varying number of Ig domains in the extracellular parts [36]. Furthermore, VEGF receptor-like molecules have also been described in invertebrates such as *Hydra* [36]. We therefore carried out additional BLASTP searches on the *E. multilocularis* genome using human VEGF- and PDGF-receptors as queries, but only obtained significant hits with the above mentioned TKDs of EmFR1, EmFR2, and EmFR3 (data not shown). These data indicated that members of the VEGF- and PDGF-receptor families are absent in *Echinococcus*, as has already been described for the closely related schistosomes [24].

Genes encoding canonical FGF ligands have so far neither been identified in genome projects of free-living flatworms [37], nor in those of trematodes [38] or cestodes [14]. In vertebrates [15] as well as several invertebrate phyla [39-41], however, canonical FGF ligands are clearly expressed. To investigate the situation more closely, we carried out BLASTP and BLASTX analyses against the predicted proteins and *E. multilocularis* contig information, respectively, using FGF ligand sequences of insect, nematode, and cnidarian origin as queries. The product of only one *E. multilocularis* gene, EmuJ_000840500 (annotated as ‘conserved hypothetical protein’) showed certain similarity to these FGF ligands and according SMART protein domain analyses could contain a FGF-ligand domain between amino acids 166 and 258, although this prediction was clearly below the prediction threshold and of low probability (E-value: 817). No export directing signal peptide was predicted for the EmuJ_000840500 protein, as would be typical for FGF ligands. Furthermore, although EmuJ_000840500 had clear orthologs in the cestodes *Taenia solium* (TsM_000953800) and *Hymenolepis microstoma* (HmN_000558500), none of these gene models had any prediction of an FGF-ligand domain in SMART analyses (nor predicted signal peptides). We thus considered it highly unlikely that EmuJ_000840500 encodes a so far not identified flatworm FGF-ligand.

Taken together, our analyses indicated that *E. multilocularis* contains genomic information for three members of the FGFR family of receptor tyrosine kinases with either one (EmFR1) or two (EmFR2, EmFR3) extracellular Ig-like domains, of which one, EmFR3, showed higher divergence within the TKD as it contained only 22 of otherwise 30 highly conserved amino acid residues of tyrosine kinases. On the other hand, no members of the VEGF- and PDGF-receptor families are encoded by the *E. multilocularis* genome, nor does it contain genes coding for canonical FGF-ligands.

### Expression of FGF receptors in *E. multilocularis* larval stages

To investigate gene expression patterns of the *Echinococcus* FGF receptors in parasite larvae, we first inspected Illumina transcriptome data for parasite primary cells after 2 and 11 days of culture (PC2d, PC11d, respectively), metacestode vesicles without or with brood capsules (MC-, MC+, respectively), as well as protoscoleces before or after activation by low pH/pepsin treatment (PS-, PS+, respectively), that had be produced during the *E. multilocularis* genome project [14]. According to these data, *emfr1* was moderately expressed in primary cells, metacestode vesicles, and protoscoleces (Fig S3). Likewise, *emfr2* was expressed in all stages, but very lowly in primary cells, somewhat more in metacestode vesicles, and highest in protoscoleces. *emfr3*, on the other hand, was low to moderately expressed in primary cells, low in metacestode vesicles, and highest in protoscoleces (Fig S3). Since primary cell preparations are characterized by a much higher content of germinative (stem) cells than metacestode vesicles [6], these expression patterns could indicate that *emfr3* is stem cell specifically expressed. We therefore carried out semi-quantitative RT-PCR experiments on cDNA preparations from *in vitro*-cultivated metacestode vesicles (MV) versus metacestode vesicles after treatment with hudroxyurea (MV-HU) or the Polo-like kinase inhibitor BI 2536 (MV-BI), in which the stem cell population had been selectively depleted [6,42]. While *emfr1* was well expressed in MV as well as MV-HU and MV-BI, both *emfr2* and *emfr3* transcripts were barely detectable in MV-HU and MV-BI (data not shown). This indicated that *emfr1* does not have a typical stem cell specific expression pattern whereas *emfr2* and *emfr3* could be expressed in the parasite’s stem cells or their immediate progeny (since HU- and BI 2536-treatment has to be carried out for at least one week [6,42]).

To clarify the situation we carried out whole-mount *in situ* hybridization experiments on metacestode vesicles according to recently established protocols [6,43,44]. In these experiments, proliferating parasite stem cells were labeled by incorporation of the nucleotide analog EdU, which was combined with detection of gene transcripts by using fluorescently labeled probes. According to these experiments, *emfr1* was expressed at low levels throughout the germinal layer and especially during the development of protoscoleces (Fig 2). The intensity of the signal was heterogenous, but no clear pattern could be discerned and it was too low to clearly determine the percentage of positive cells.

**Fig 2.**
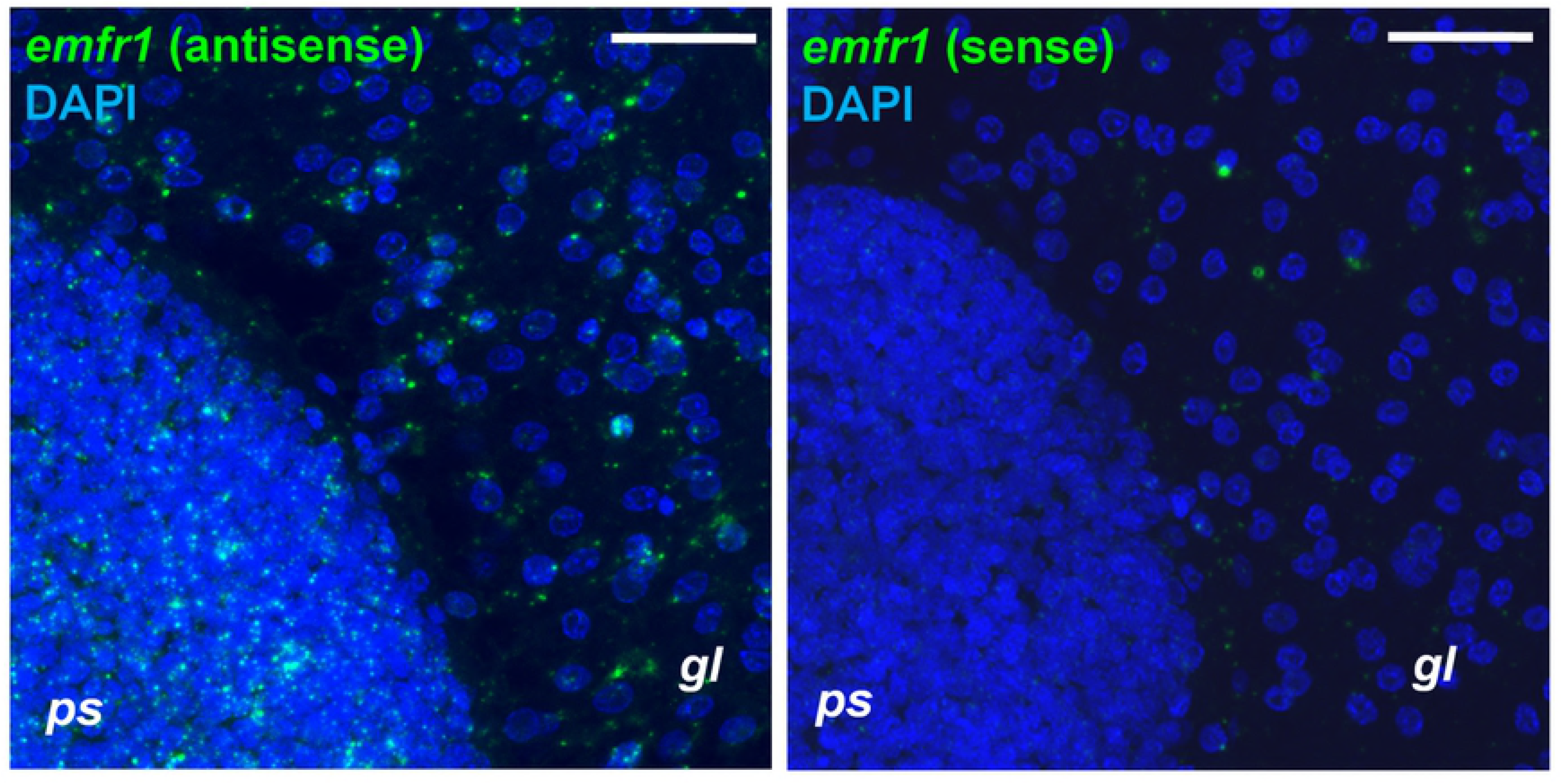
WMISH analysis of *emfr1* expression. Left panel, antisense probe detecting *emfr1* expression. Right panel, sense probe (negative control). WMSIH signal is shown in green, nuclear DAPI staining in <u>blue. *gl*</u>, germinal layer; *ps*, developing protoscolex. Bar: 20 µm

*emfr2*, on the other hand, was specifically expressed in a population of small sized cells, which comprised 1.7% to 6.3% of all cells in the germinal layer (Fig 3A). None of these *emfr2*^+^ cells were also EdU^+^, indicating that they were post-mitotic [6]. During initial protoscolex development, *emfr2*^+^ cells accumulated in the peripheral-most layer of cells, as well as in the anterior-most region (which will form the rostellum), and in three longitudinal bands of cells in the interior of the protoscolex buds (Fig 3B). Again, practically none of the *emfr2* labelled cells was EdU^+^ (less than 1% of *emfr2*^+^ cells were EdU^+^; Fig 3C). Later during development, some *emfr2*^+^ cells were found in the protoscolex body, but most accumulated in the developing rostellum and the suckers (Fig 3D). Importantly, *emfr2* expression was restricted to the sucker cup, where cells are differentiating into *em-tpm-hmw*^+^ muscle cells [6], and not the sucker base, where EdU^+^ germinative cells accumulate [6] (Fig 3D). Taken together, these data indicate that *emfr2* is expressed in post-mitotic cells, of which many are likely to be differentiating or differentiated muscle cells.

**Fig 3.**
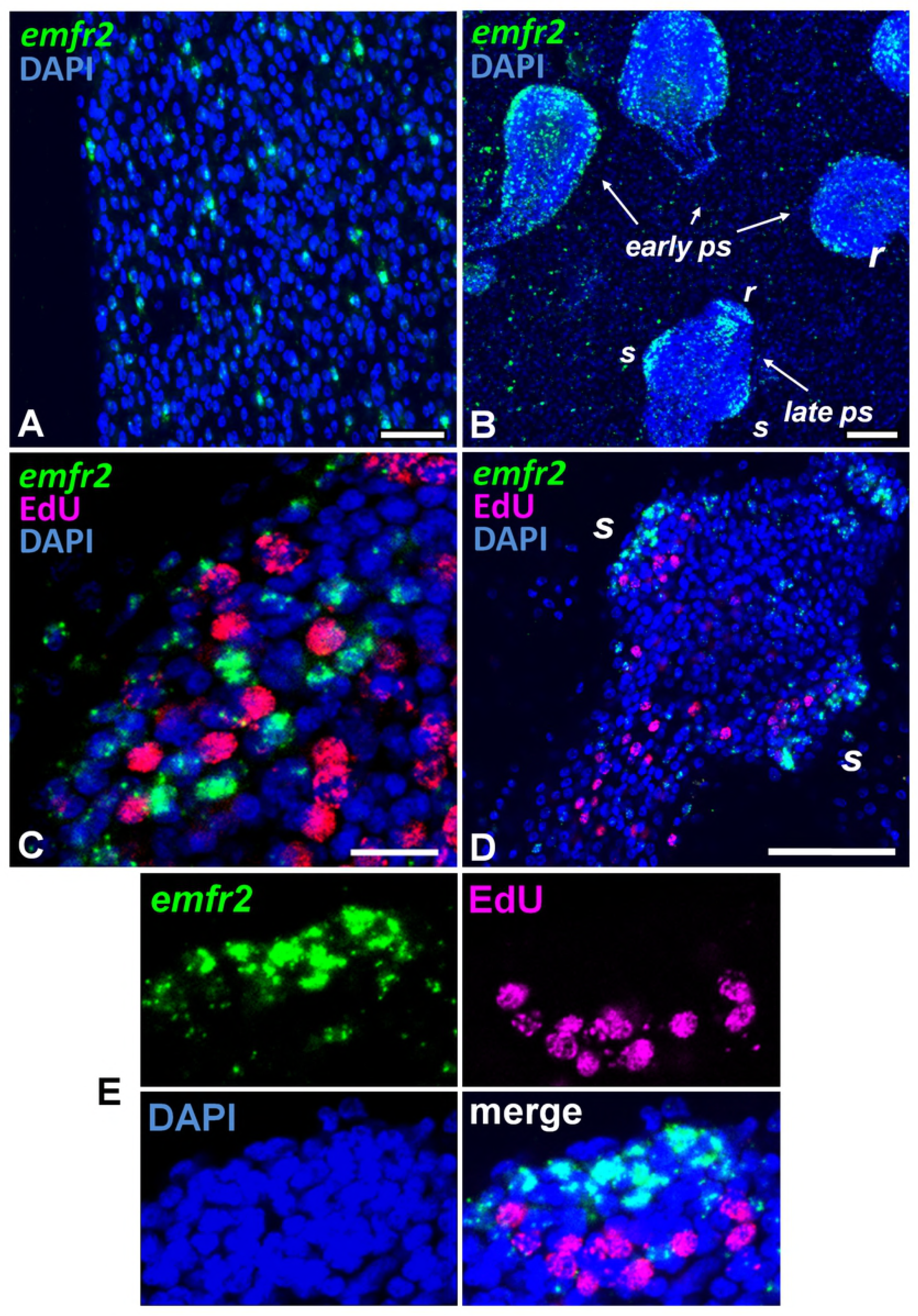
WMISH analysis of *emfr2* expression. Shown are WMISH analyses of *emfr2* expression (A, B) and co-detection of *emfr2* WMISH and EdU incorporation (C, D, E) during metacestode development. In C, D, and E, metacestodes were cultured in vitro with 50 µM EdU for 5 h and were then processed for WMISH and EdU detection. In all panels the WMISH signal is shown in green, EdU detection in red, and DAPI nuclear staining in blue. A, detail of the germinal layer. B, general view of a region of a metacestode with protoscoleces in different stages of development. C, early protoscolex development. D, late protoscolex development. E, Detail of a sucker, maximum intensity projection of a confocal Z-stack. *r*, rostellum; *s*, sucker. Bars are 20 µm for A, 50 µm for B, 10 µm for C, and 50 µm for D. Sense probe (negative control) was negative in all samples.

Finally, *emfr3* was expressed in very few cells in the germinal layer (less than 1% of all cells, although the number is difficult to estimate since they were absent in most random microscopy fields) (Fig 4A). *emfr3*^+^ cells accumulated in small numbers during brood capsule and protoscolex development (Fig 4B). The *emfr3*^+^ cells had a large nucleus and nucleolus and, thus, had the typical morphology of germinative cells (Fig S4). At the final stages of protoscolex development, few *emfr3*^+^ cells were present in the body region and some signals were also present in the rostellar region (Fig S4). In the developing protoscolex and in the germinal layer, *emfr3*^+^ EdU^+^ double positive cells were found (Fig 4). These data indicated that *emfr3* is expressed in a very small number of proliferating cells with the typical morphology of germinative cells.

**Fig 4.**
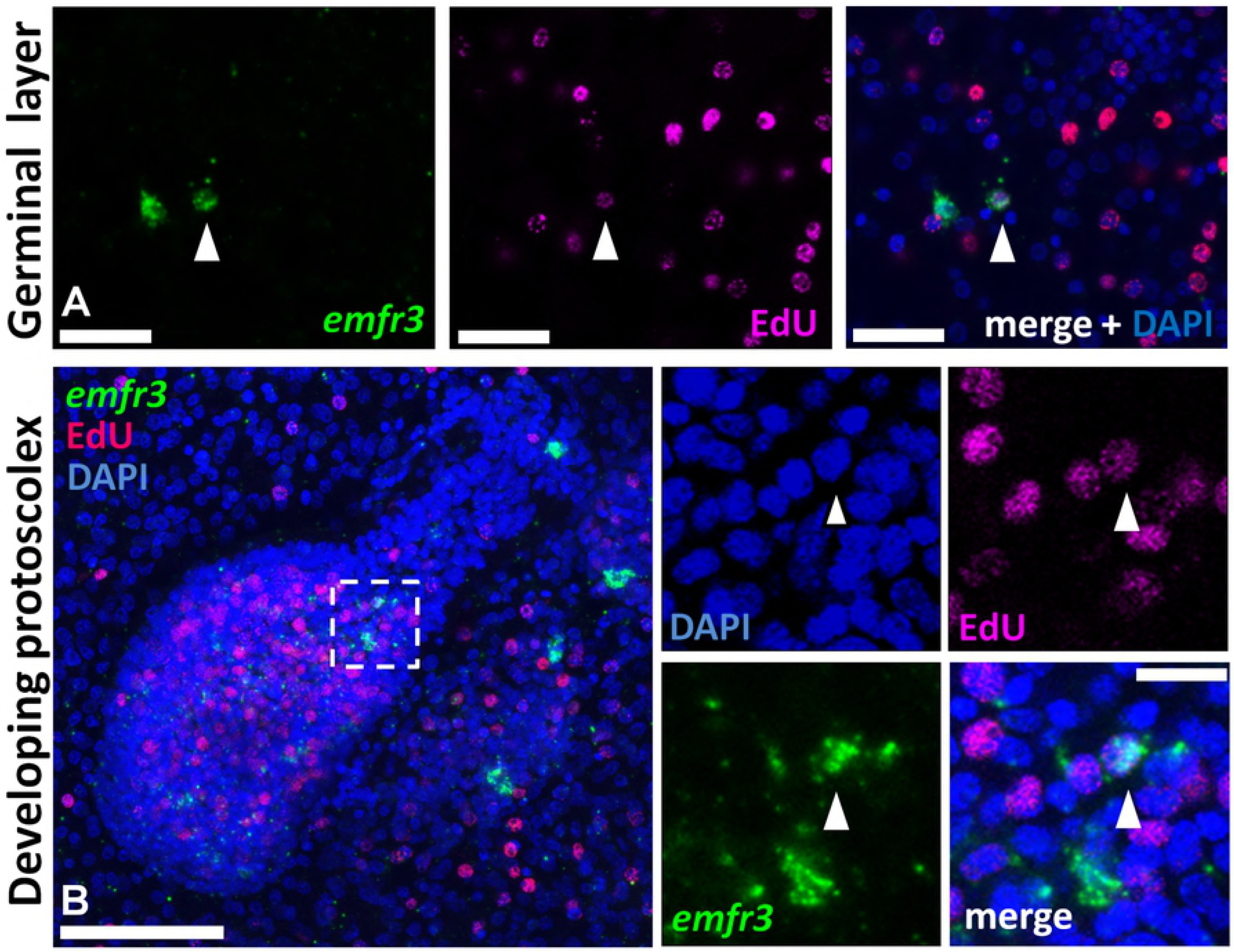
Co-detection of *emfr3* WMISH and EdU incorporation during metacestode development. Metacestodes were cultured in vitro with 50 µM EdU for 5 h and were then processed for WMISH and EdU detection. In panels displaying multiple channels, the WMISH signal is shown in green, EdU detection in red, and nuclear DAPI staining in blue. A, germinal layer. B, developing protoscolex. Arrowheads indicate Edu^+^ *emfr3*^+^ double-positive cells. Bars are 25 µm for A, 50 µm for B (main panel) and 10 µm for B (enlarged inset).

In summary, the three *E. multilocularis* FGF receptor genes showed very different expression patterns in metacestode and protoscolex larval stages. While *emfr1* was lowly expressed in cells that are dispersed throughout the germinal layer, *emfr2* displayed an expression pattern indicative of differentiating/differentiated muscle cells. *emfr3*, on the other hand, appeared to be expressed in a minor sub-population of germinative cells.

### Host FGF stimulates *E. multilocularis* larval proliferation and development

Since *E. multilocularis* larvae do not express canonical FGF ligands (see above), but usually develop in an environment in which FGF1 and FGF2 are abundant [17], we next tested whether host-derived FGF ligands can stimulate parasite development *in vitro*. To this end, we employed two different cultivation systems which we had previously established [7-10]. In the axenic metacestode vesicle cultivation system, mature metacestode vesicles of the parasite are cultivated in the absence of host cells under reducing culture conditions (e.g. low oxygen)[7,9]. In the primary cell cultivation system [8,10], axenically cultivated metacestode vesicles are digested to set up cell cultures which are highly enriched in parasite germinative (stem) cells (~ 80% [6]), but which also contain certain amounts of differentiated cells such as muscle or nerve cells. In the primary cell cultivation system, mature metacestode vesicles are typically formed within 2 – 3 weeks, which is critically dependent on proliferation and differentiation of the germinative cell population [6] in a manner highly reminiscent of the oncosphere-metacestode transition [3].

As shown in Fig 5A, the exogenous addition of 10 nM FGF1 to mature metacestode vesicles already resulted in an elevated incorporation of BrdU, indicating enhanced proliferation of parasite stem cells, which was even more pronounced in the presence of 100 nM FGF1. In the case of FGF2, addition of 100 nM resulted in enhanced BrdU incorporation in a statistically significant manner (Fig 5A). Likewise, metacestode vesicles cultured for 4 weeks in the presence of 10 nM FGF1 or FGF2 displayed a considerably larger volume (about two-fold) when compared to non-FGF-stimulated vesicles (Fig 5B).

**Fig 5.**
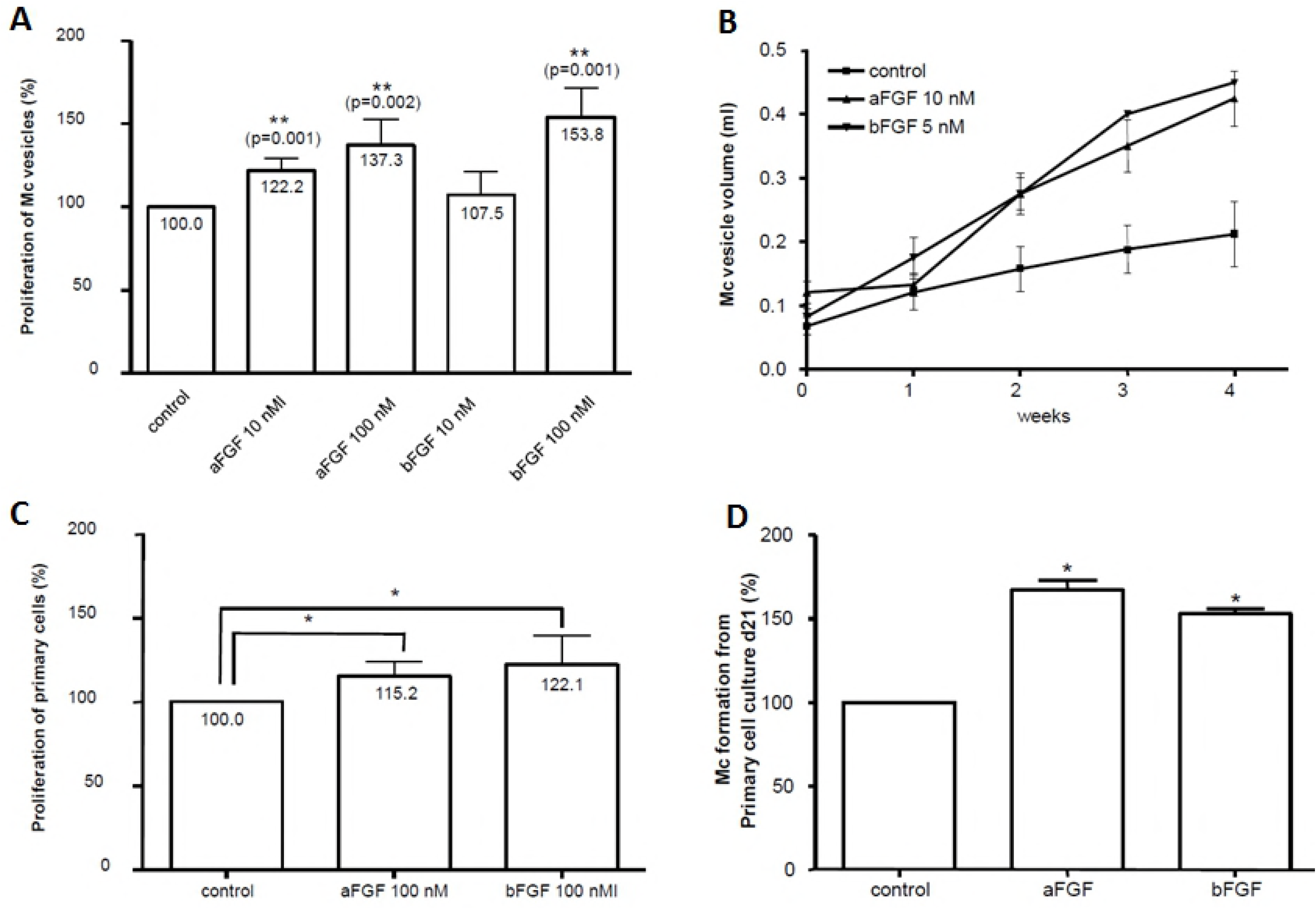
Effects of host FGFs on *E. multilocularis* proliferation and development. **A**, effects of FGFs on metacestode proliferation. Axenically cultivated metacestode vesicles (eight per well) were incubated for 48 h in DMEM medium (without FCS), then BrdU (1 mM) and FGFs (10 or 100 nM) were added and incubation proceeded for another 48 h before cell lysis, DNA isolation, and BrdU detection. Bars represent percentage of BrdU incorporation with the cMEM control set to 100%. Statistical evaluation of four independent experiments (n=4) which were conducted in duplicates is shown. Student’s t-test (two tailed): *p<0.05. **B**, effects of host FGFs on metacestode vesicle development. Single axenically generated metacestode vesicles were *in vitro* cultivated for four weeks in the presence of 10 nM FGF1 (aFGF) or 5 nM FGF2 (bFGF) with daily medium changes. Control vesicles were kept in cMEM. Growth (in ml) of vesicles was monitored. In each of three independent experiments (n=3), four vesicles were examined for every single condition. **C**, effects of FGFs on *E. multilocularis* primary cell proliferation. Freshly isolated *E. multilocularis* primary cells were incubated for 48 h in 2 ml cMEM, then BrdU (1 mM) and FGFs (100 nM) were added and cells were incubated for another 48 h before DNA isolation and BrdU detection. Statistical analysis was performed as in A. D, effects of host FGFs on metacestode vesicle development. Freshly prepared E. multilocularis primary cells were cultured for 21 days in cMEM medium in the presence or absence of 100 nM FGF1 (aFGF) or FGF2 (bFGF). Half of the medium volume was renewed every second day. The number of newly formed metacestode vesicles at day 21 was analysed. The bars represent the percentage of formed vesicles with the cMEM control set to 100%. The statistical evaluation of three independent experiments (n=3) which were conducted in duplicates is shown. Student’s t-test (two-tailed): *p<0.05.

In the primary cell cultivation system, 100 nM concentrations of host ligands had to be added to observe statistically significant effects. Again, the incorporation of BrdU by primary cell cultures was stimulated in the presence of host-derived FGF ligands (Fig 5C), as was the formation of mature metacestode vesicles from primary cell cultures (Fig 5D).

Taken together, these results indicated that host-derived FGF ligands, and in particular FGF1, can stimulate cell proliferation and development of *E. multilocularis* primary cell cultures and mature metacestode vesicles.

### The *E. multilocularis* FGF receptors are activated by host-derived FGF ligands

Having shown that host-derived FGF ligands can stimulate parasite proliferation and development *in vitro*, we were interested whether these effects might be mediated by one or all three of the parasite’s FGF receptors which are expressed by the metacestode larval stage. To this end, we first made use of the *Xenopus* oocyte expression system in which the activity of heterologously expressed protein kinases can be measured by germinal vesicle breakdown (GVBD). This system has previously been used to measure the activities of the TKD of schistosome FGF receptors [24], as well as the host-EGF (epidermal growth factor) dependent activation of a schistosome member of the EGF receptor family [31]. We, thus, expressed *Pleurodeles* FGFR1 (as a positive control), which is highly similar to human FGFR1 [32], and all three parasite FGF receptors in *Xenopus* oocytes, which were then stimulated by the addition of 10 nM FGF1 or FGF2. As negative controls, we also expressed kinase-dead versions of all *Echinococcus* FGF receptors in *Xenopus* oocytes.

As can be deduced from Table 1, control (non-stimulated) oocytes were negative for GVBD, whereas progesterone-stimulated oocytes displayed 100% vesicle breakdown. Expression of FGFR1 did not yield GVBD but, after stimulation with 10 nM FGF1, 100% of oocytes underwent GVBD, indicating stimulation of the *Pleurodeles* receptor by FGF1 (as expected). Upon expression of any of the parasite receptors in *Xenopus* oocytes, no GVBD was observed when no ligand was added. The addition of 10 nM FGF1 to these receptors, however, resulted in 100% GVBD for EmFR1 and EmFR2, as well as to 80% GVBD in the case of EmFR3. In the case of human FGF2 (10 nM), 90% GVBD was observed for EmFR1 and EmFR3, and 85% for EmFR2. No GVBD was observed upon addition of 10 nM FGF1 or FGF2 when the kinase-dead versions of the parasite receptors were expressed (data not shown and Fig S5). These data clearly indicated that all three parasite receptors were responsive to host derived FGF ligands (albeit to somewhat different extent) and that the kinase activity of the parasite receptors was essential to transmit the signal. We also measured the phosphorylation state of the parasite FGF receptors upon addition of exogenous FGF1 and FGF2 (10 nM each) to *Xenopus* oocytes and obtained significantly induced levels of tyrosine phosphorylation in all three cases (Fig S5).

**Table 1.** Effects of host FGFs and BIBF 1120 on *E. multilocularis* FGF receptors. **Table 1.** *Pleurodeles* FGFR1 (FGFR1) and the three *E. multilocularis* FGF receptors (EmFR1, EmFR2, EmFR3, as indicated) were expressed in *Xenopus* oocytes without or with human FGF1 (hFGF1) or FGF2 (hFGF2) and in the presence of different concentrations of BIBF 1120 (as indicated). Germinal vesicle breakdown (GVBD) was monitored after 15 h of incubation. Numbers indicate % of GVBD, mean of two independent experiments.

| BIBF1120 |  | 1μM | 10μM | 20μM |
| --- | --- | --- | --- | --- |
| control | 0 | 0 | 0 | 0 |
| Progesterone | 100 | 100 | 100 | 70 |
| FGFR1 | 0 | 0 | 0 | 0 |
| FGFR1 + hFGF1 | 100 | 40 | 0 | 0 |
| EmFR1 | 0 | 0 | 0 | 0 |
| EmFR1 + hFGF1 | 100 | 90 | 0 | 0 |
| EmFR1 + hFGF2 | 90 | 50 | 0 | 0 |
| EmFR2 | 0 | 0 | 0 | 0 |
| EmFR2 + hFGF1 | 100 | 50 | 0 | 0 |
| EmFR2 + hFGF2 | 85 | 0 | 0 | 0 |
| EmFR3 | 0 | 0 | 0 | 0 |
| EmFR3 + hFGF1 | 80 | 80 | 30 | 0 |
| EmFR3 + hFGF2 | 90 | 80 | 30 | 0 |

Taken together, these data indicated that all three parasite FGF receptors were activated by binding of host-derived FGF1 and FGF2, which was followed by auto-phosphorylation of the intracellular kinase domain and downstream transmission of the signal to the *Xenopus* oocyte signaling systems which induce GVBD.

### Inhibition of the *E. multilocularis* FGF receptors by BIBF 1120

The small molecule compound BIBF 1120 (also known as Nintedanib or Vargatef™) is a well-studied and highly selective, ATP-competitive inhibitor of mammalian members of the FGF-, VEGF-, and PDGF-receptor families with very limited affinity to other receptor tyrosine kinases [45,46]. As a possible agent to selectively inhibit FGF receptor tyrosine kinase activities in the parasite, we measured the effects of BIBF 1120 on EmFR1, EmFR2, and EmFR3 upon expression in the *Xenopus* oocyte system. As can be deduced from Table 1, a concentration of 1 µM of exogenously added BIBF 1120 already diminished the activity (after stimulation with 10 nM FGF1) of FGFR1 to 40%, and led to a complete block of kinase activity upon addition of 10 µM BIBF 1120. In the case of EmFR1 and EmFR2, 1 µM BIBF 1120 also led to a marked decrease of receptor kinase activity, although to a somewhat lower extent than in the case of FGFR1. In the presence of 10 µM BIBF 1120, on the other hand, the activities of EmFR1 and EmFR2 were completely blocked. In the case of EmFR3, exogenous addition of 1 µM BIBF 1120 had only slight effects on GVBD, whereas 10 µM BIBF1120 reduced the activity to less than 50% and 20 µM BIBF 1120 completely blocked tyrosine kinase-dependent GVBD (Table 1). Upon addition of 20 µM BIBF 1120, tyrosine kinase activity of all receptors tested was completely inhibited.

Taken together, these data indicated that all three *Echinococcus* FGF receptor tyrosine kinases were affected by BIBF 1120, although in all three cases higher concentrations of the inhibitor were necessary to completely block tyrosine kinase activity when compared to FGFR1. BIBF 1120 treatment had the lowest effects on the activity of EmFR3.

### BIBF 1120 inhibits *E. multilocularis* larval development

We next tested the effects of different concentrations of BIBF 1120 on parasite development and survival in the primary cell and metacestode vesicle culture systems. As shown in Fig 6A, the addition of 1 – 10 µM BIBF 1120 had clear concentration dependent effects on mature metacestode vesicle survival which, after cultivation for 18 days, led to about 20% surviving vesicles in the presence of 1 µM BIBF 1120, 10% surviving vesicles in the presence of 5 µM BIBF 1120, and no survival when 10 µM BIBF 1120 was applied. To test whether the metacestode vesicles were indeed no longer capable of parasite tissue regeneration, we set up primary cell cultures from microscopically intact vesicles which had been treated with 5 µM BIBF 1120 for 5 days (90% intact vesicles) and let the cultures recover in medium without inhibitor. From these cultures, however, we never obtained the formation of mature vesicles (data not shown), indicating that either the parasite’s stem cell population, and/or other cell types necessary for parasite development in the primary cell culture system, were severely damaged after BIBF 1120 treatment.

**Fig 6.**
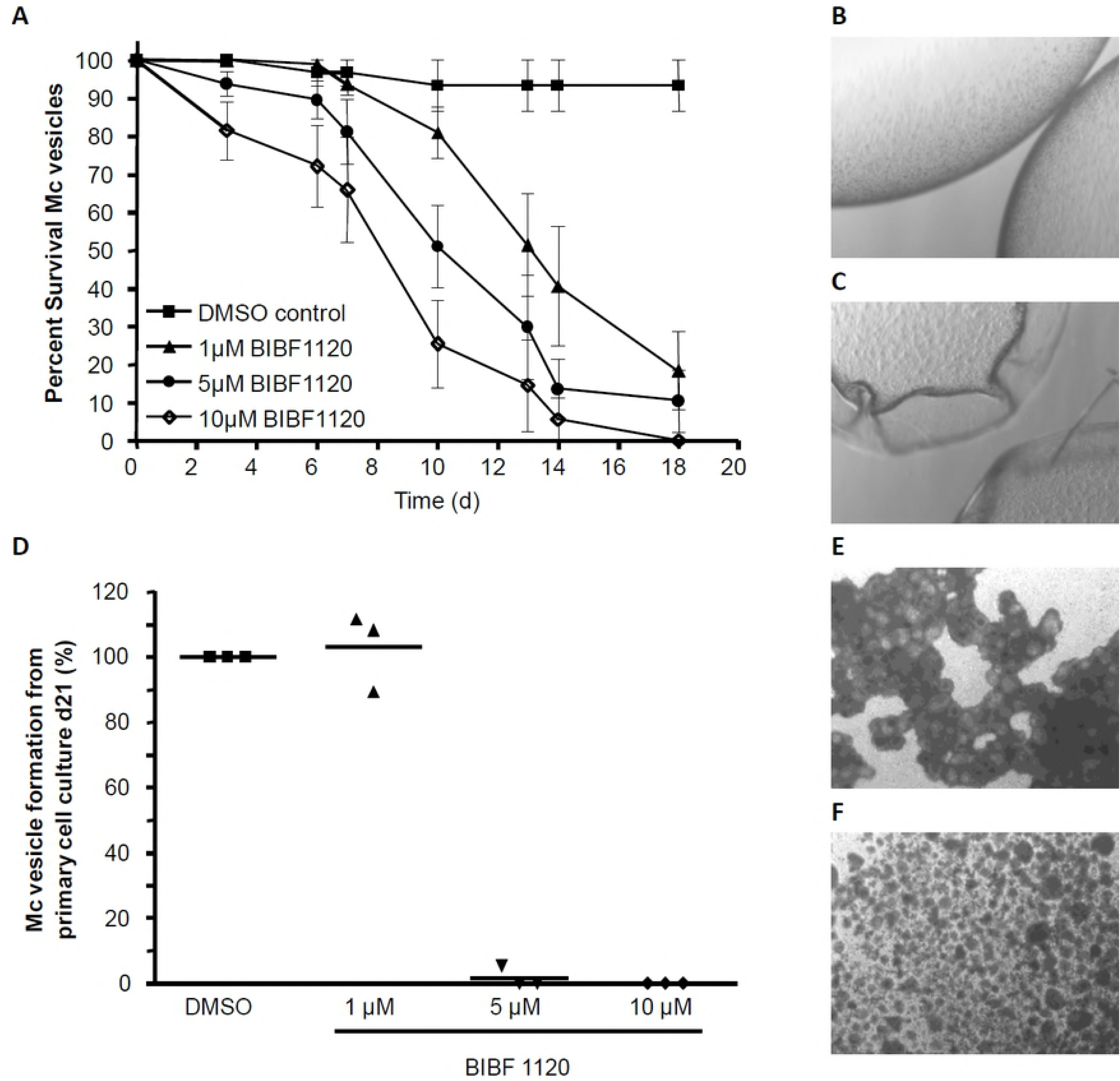
Effects of BIBF 1120 on *E. multilcoularis* metacestode vesicles and primary cells. A, axenically cultivated metacestode vesicles (8 per well in 2 ml volume) were incubated for 18 days in the presence of 0.1 % DMSO or BIBF 1120 in different concentrations (as indicated) and the structural integrity of the vesicles was monitored. Structurally intact vesicles (B) and damaged vesicles (C) are shown to the right. The experiment was repeated three times in duplicates. D, freshly isolated *E. multilocularis* primary cells were cultured for 21 days in cMEM medium in the presence of DMSO (0.1 %) or BIBF 1120 at different concentrations (as indicated). After 21 days, newly formed metacestode vesicles were counted. The DMSO control for each of the three independent experiments was set to 100%. Microscopic images (25x magnification) of cultures after 2 weeks with DMSO (E) or 10 µM BIBF 1120 (F) are shown to the right.

We then also tested the effects of BIBF 1120 on fresh primary cell cultures from previously untreated metacestode vesicles. As shown in Fig 6B, a concentration of 1 µM BIBF 1120 had no effect on the formation of metacestode vesicles from these cultures, whereas vesicle formation was completely blocked in the presence of 5 µM or 10 µM BIBF 1120.

Altogether, these results clearly indicated detrimental effects of BIBF 1120 on parasite development already at concentrations as low as 5 µM. Since the parasite does not express known alternative targets for BIBF 1120, such as VEGF- or PDGFR-receptors, we deduced that these effects are due to the inhibition of one or more of the parasite’s FGF receptor tyrosine kinases.

### Host FGF ligands stimulate Erk signaling in *E. multiocularis*

One of the major downstream targets of FGF signaling in other organisms is the Erk-like MAPK cascade, a complete module of which we had previously identified in *E. multilocularis* [33,47,48]. In particular, we had previously shown that the phosphorylation status of the parasite’s Erk-like MAP kinase, EmMPK1, can be measured by using antibodies against the phosphorylated form of the human Erk-kinase [33]. To investigate whether exogenously added host FGF can affect the *E. multilocularis* Erk-like MAPK cascade module, we first incubated mature metacestode vesicles for 4 days in serum-free medium (which has no effect on vesicle integrity [29]) and then stimulated these vesicles for 30 sec, 60 sec, and 60 min with 10 nM FGF1 and FGF2. As shown in Fig 7A, FGF1 treatment had a clear effect of EmMPK1 phosphorylation already after 30 sec of exposure. In the case of FGF2, the effect was still measurable, but clearly less pronounced than in the case of FGF1 (Fig. 7A). We then also measured the effects of BIBF 1120 treatment on EmMPK1 phosphorylation. To this end, metacestode vesicles were incubated in hepatocyte-conditioned medium and were then subjected to BIBF 1120 treatment (5 µM, 10 µM) for 30 min. As shown in Fig 7B, this led to diminished phosphorylation of EmMPK1 when 10 µM BIBF 1120 was used.

Taken together, these data indicated that, like in mammals and other invertebrates, the *E. multilocularis* Erk-like MAPK cascade can be activated through FGF signaling, initiated by exogenously added, host-derived FGF ligands.

**Fig 7.**
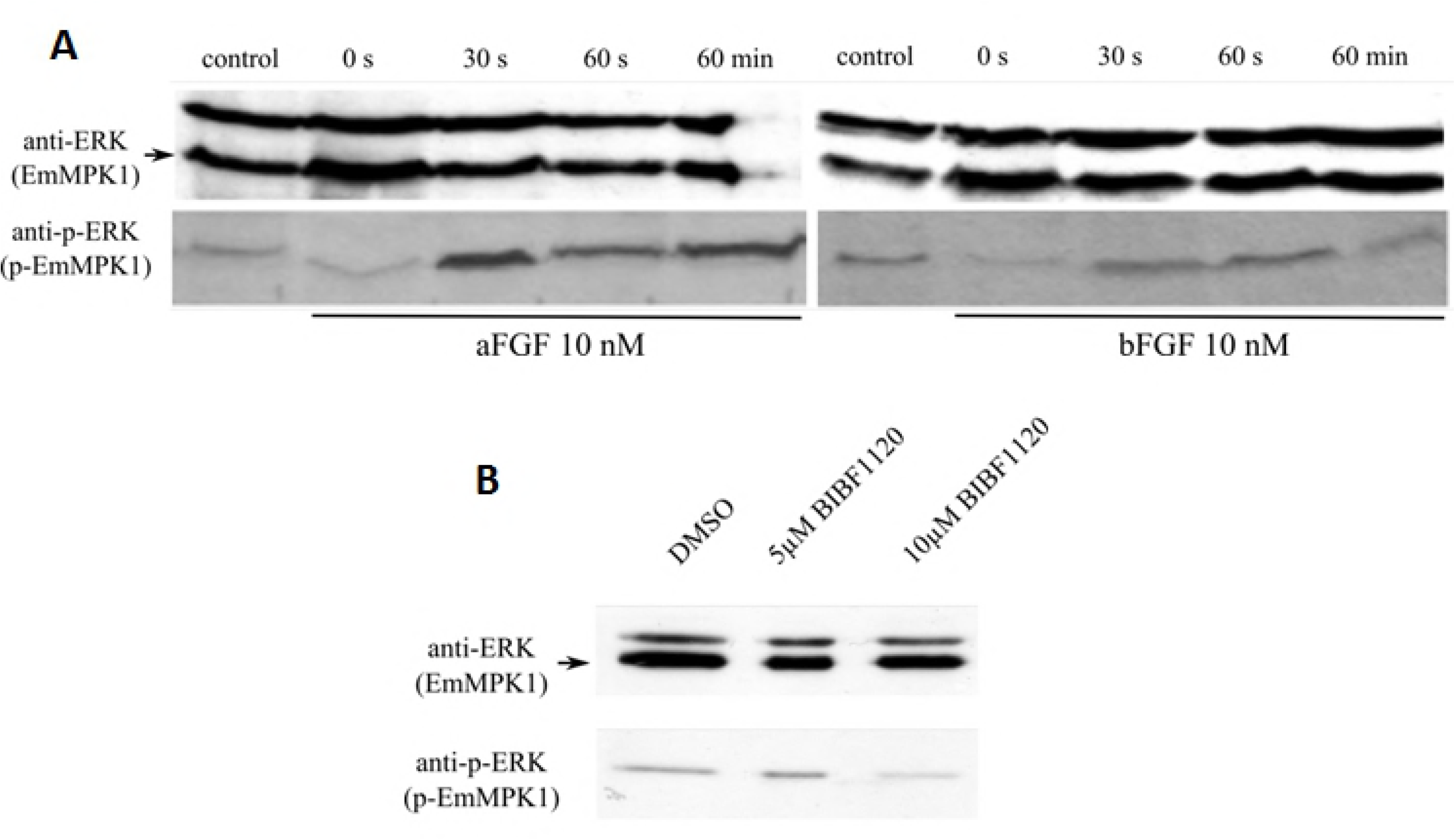
Effects of host FGFs and BIBF 1120 on EmMPK1 phosphorylation in metacestode vesicles. **A**, axenically cultivated metacestode vesicles were incubated in cMEM (control) or in medium without FCS (0 s), upon which FGF1 (aFGF) or FGF2 (bFGF) were added at a concentration of 10 nM for 30 sec (30s), 60 sec (60s) or 60 min (60min). Protein lysates were subsequently separated by 12% SDS-PAGE and Western blots were analysed using polyclonal antibodies against Erk-like MAP kinases (anti-ERK) or double phosphorylated Erk-like MAP kinases (anti-p-ERK). **B**, axenically cultivated metacestode vesicles were incubated with DMSO (negative control), 5 mM or 10 mM BIBF1120 (30 min each) and cell lysates were subsequently analysed as described above. Both experiments were performed in triplicate with similar results.

## Discussion

An important difference between parasitic helminths and all other infectious agents (excluding viruses) is that these organisms are evolutionarily relatively closely related to their vertebrate or invertebrate hosts with which they share an ancestor that has lived around 500 to 600 million years ago. Since all metazoans share evolutionarily conserved signalling systems for cell-cell communication, this opens the possibility for host-parasite cross-communication involving evolutionarily conserved cytokines of one partner (e.g. the host) and cognate receptors of the other (e.g. the parasite), which would be of particular relevance for systemic helminths that infect host organs. Several previous studies indicated that this type of host-pathogen interaction is indeed important in helminth infections. In the *E. multilocularis* system, which develops in close association with host liver tissue, we previously demonstrated that host insulin stimulates parasite development and growth via acting on evolutionarily conserved insulin-receptor tyrosine kinases that are expressed by the parasite [11]. A similar type of cytokine receptor interaction appears to involve epidermal growth factor (EGF)-like cytokines and cognate parasite receptors, of which three (EmER, EmERb, EmERc) are expressed by *E. multilocularis* larvae [3,12]. As recently shown by Cheng et al. [13], host-derived EGF is able to stimulate germinative cell proliferation in *in vitro* cultivated parasite larvae and can stimulate at least one of the parasite EGF-receptors, EmER, when heterologously expressed in *Xenopus* oocytes. Although the parasite itself expresses several EGF-like molecules [14], which likely act on its EGF receptors, these data indicate that host-EGF could act as an additional stimulus, particularly in response to liver tissue damage as it is inflicted upon entry of the parasite into the host liver [4]. Apart from insulin- and EGF-signalling systems, host-parasite interactions in larval cestode infections might also involve the family of transforming growth factor (TGF)-β/bone morphogenetic protein (BMP)-family of cytokines since host TGF-β has very recently been shown to stimulate larval growth of the cestode *Taenia crassiceps in vitro* and was found to interact with parasite TGF-β receptors [49]. In a similar way, human BMP2 was reported by us to interact with an *E. multilocularis* BMP receptor [50], although no direct effects of host BMP on parasite development were yet observed. Like in cestodes, the occurrence of insulin- and EGF-receptor tyrosine kinases as well as TGF-β/BMP serine/threonine kinases with the capability of interacting with respective human hormones/cytokines was reported for schistosomes [31,51-55], and stimulatory effects of host EGF on the development of schistosome snail stages were observed [31]. In the present work, we extend the list of respective host-parasite cross-communication systems to the FGF-family of host cytokines and cognate FGF receptor tyrosine kinases which, to our knowledge, have never before been addressed in this context.

We herein clearly show that mammalian FGF1 and FGF2 stimulate the development of *E. multilcularis* metacestode vesicles from cultivated parasite primary cells, which are highly enriched in germinative (stem) cells [6]. Furthermore, we show that human FGF also stimulates proliferation and growth of mature metacestode vesicles. Both FGF1 and FGF2 are abundantly present in mammalian liver tissue where they are mostly released upon liver cell damage and during regeneration processes [17-20]. Although the precise amounts of host FGF in periparasitic lesions of *E. multilocularis* infected mice has not been measured to date, it is very likely that the early establishing metacestode is encountering considerable amounts of these cytokines since extensive damage to liver tissue is observed not only in chronically infected mice but also in early stages of the infection [4]. The early infectious stage is critical in the establishment of the parasite since the invading oncosophere is not yet producing the laminated layer (LL), an important structure that protects the parasite from direct actions of immune cells [56]. The laminated layer surrounds mature metacestode vesicles (established after 1-2 weeks after invasion) in the chronic phase of the disease. The stimulation of parasite development from stem cells towards mature metacestode vesicles by host FGF, as observed in our cultures, could thus help the parasite to abridge this critical phase and to successfully establish within the liver. Chronic AE is characterized by extensive liver fibrosis, particularly in the peri-parasitic area [57] and is thought to be mediated by hepatic stellate cells [5], which greatly upregulate FGF release during liver regeneration and fibrosis induction [20]. Hence, not only in the initial phase of parasite establishment, but also in the chronic phase of AE, the *E. multilocularis* metacestode should be in contact with elevated levels of host FGF that can continuously support parasite growth and proliferation.

Using the *Xenopus* expression system we clearly showed that all three identified FGF receptors of *E. multilocularis* are functionally active kinases that are capable of inducing GVBD when properly stimulated. We also demonstrated that all three *Echincoccus* kinases are responsive to human FGF1 and FGF2, albeit to somewhat different extent. While 10 nM FGF1 fully stimulated both EmFR1 and EmFR2, EmFR3 was less activated than the other receptors by both FGFs. This does not necessarily indicate, however, that human FGFs bind less well to EmFR3 than to the other two receptors. Instead, EmFR3 might be activated to a lesser extent since two tyrosine residues, Y653 and Y654, which in human FGFR1 are necessary for full activation of the receptor [35], are conserved in EmFR1 and EmFR2 but absent in EmFR3 (Fig S2). In any case, our data clearly show that particularly human FGF1, but also human FGF2, are capable of activating the parasite receptors. Since the parasite apparently does not produce intrinsic FGF ligands, the only canonical FGFs it encounters during liver invasion are host derived. It is, thus, logical to assume that the effects of FGF1 and FGF2 on parasite growth and development are mediated by one, two or all three *Echinococcus* FGF receptors. In line with this assumption is the stimulation of the parasite’s Erk-like MAPK cascade module upon exogenous addition of host FGF to metacestode vesicles. In mammals, the two prominent downstream signalling pathways of FGF receptors are the Ras-Raf-MAPK cascade and the PI3K/Akt pathway, while two others are STAT signalling and the phospholipase γ (PLCγ) pathway [15]. The STAT signalling pathway is absent in *Echinococcus* [14] and PLCγ involves binding to human FGF receptors at tyrosine residues that are not conserved in the *Echinococcus* receptors. We had previously demonstrated that the PI3K/Akt pathway exists in *Echinococcus* [11] but we could not measure differential phosphorylation of EmAKT in response to exogenous FGF or after metacestode vesicle treatment with BIBF 1120 (data not shown), indicating that in contrast to human cells, *Echinococcus* mainly involves the MAPK cascade pathway for downstream FGF signalling.

Concerning cellular expression patterns we detected significant differences between the *Echincoccus* FGF receptors and those of the related schistosomes and planaria. In planaria, the expression of FGF receptors is a hallmark of neoblast stem cells and also occurs in cephalic ganglions [21,23]. In schistosomes, one of the two FGF receptors, *fgfr*A, is a marker for a prominent subset of somatic stem cells [25-27] and, like *fgfr*B, is also expressed in the reproductive organs [24]. In *Echinococcus*, we found only one receptor, EmFR3, which is expressed in germinative cells, but only in a very tiny subpopulation. EmFR2 was also only expressed in a few cells which were, however, clearly post-mitotic and most probably represented muscle cells. Likewise, the third *Echinococcus* FGF receptor, EmFR1, was expressed throughout the parasite’s metacestode tissue without specific association with the germinative cells. In qRT-PCR analyses on metacestode ’vesicles which were specifically deprived of stem cells after treatment with hydroxyurea [6] or the Polo-like kinase inhibitor Bi 2536 [42] we also never observed diminished levels of *emfr1* expression (data not shown), which further supports that the gene is not exclusively expressed in germinative cells. Taken together, these data indicate that the close association of FGF receptor expression with stem cells as observed in planaria and schistosomes is at least highly modified in the *Echinococcus* system, in which at least EmFR1 and EmFR2 are clearly expressed in post-mitotic cells. This adds to stem cell-specific gene expression differences which we had previously observed between planaria and cestodes e.g. concerning *argonaute* or histone deacetylase-orthologs [6], and also indicates clear differences between stem cell – specific gene expression patterns in the parasitic flatworm lineages of trematodes and cestodes.

We observed clear inhibitory effects of the tyrosine kinase inhibitor BIBF 1120 on the enzymatic activity of all three *Echinococcus* FGF receptors upon heterologous expression in the *Xenopus* system and also demonstrated that this compound can profoundly affect parasite survival and development *in vitro*. Since in the *Xenopus* system concentrations of 20 µM BIBF 1120 fully inhibited all three *Echinococcus* FGF receptors we cannot clearly state whether the detrimental effects on parasite development were due to the specific inhibition of one of the three *Echinococcus* FGF receptors, or on combined activities against all three enzymes. However, based on the fact that EmFR3 is only expressed in less than 1% of the cells of the metacestode, that *emfr3* expressing cells only accumulate during the formation of protoscoleces, and that EmFR3 showed lowest levels of inhibition upon expression in *Xenopus* cells, we do not consider this receptor as a prominent candidate for mediating the BIBF 1120 effects on primary cells and the metacestode. We also consider it unlikely that the inhibition of EmFR2 had produced these effects since the respective gene is only expressed in a small subset (2-5%) of all metacestode cells. Based on the expression of *emfr2* in muscle cells or muscle cell progenitors, however, we cannot completely rule out that EmFR2 inhibition might have led to the depletion of specific parasite cells that are necessary to form a stem cell niche for the parasite’s germinative cells. At least in planaria it has already been shown that muscle cells provide important positional information on the stem cell population [58] and our recent studies on the *wnt* signalling pathway in *Echinococcus* clearly demonstrated that this is also the case for cestodes [44]. However, based on the fact that *emfr1* is expressed throughout the metacestode and that in both primary cells and metacestode vesicles *emfr1* is the highest expressed FGF receptor gene we propose that most of the effects of BIBF 1120 on *Echinococcus* development are mediated by EmFR1 inhibition. For the development of novel chemotherapeutics against AE, EmFR1 and EmFR2 would thus be highly interesting target molecules although, of course, BIBF 1120 as originally developed against human FGF receptors showed somewhat higher activities against FGFR1 in the *Xenopus* expression system than against the parasite FGF receptors. Nevertheless, and using the activity assays developed in this work, it should be possible to identify compounds which are related to BIBF 1120 but which show higher affinities against the parasite enzymes than against human FGF receptors.

## Conclusions

In the present work we provide clear evidence that human FGF ligands are capable of activating evolutionarily conserved tyrosine kinases of the FGF receptor family that are expressed by the larval stage of *E. multilocularis* and that the parasite’s Erk-like MAPK cascade is stimulated upon exogenous addition of human FGFs to metacestode vesicles. We also showed that human FGF1 and FGF2 are stimulating the development of metacestode vesicles from parasite primary cell cultures and that they accelerate metacestode vesicle proliferation and growth in vitro. Since FGF1 and FGF2 are expressed in considerable amounts within the host liver, the primary target organ for the establishment of the *E. multilocularis* metacestode, and since FGF ligands are also constantly produced during liver regeneration and fibrosis, which are consequences of parasite growth within the intermediate host, we consider the observed *in vitro* effects on parasite FGF signalling and metacestode development also of high relevance *in vivo*. Liver-specific activities of host FGF could thus support the development of metacestode vesicles from stem cells that are delivered to the liver by the oncosphere larva, and could constantly stimulate asexual proliferation of the metacestode during an infection. We finally showed that at least one compound that inhibits the activities of mammalian FGF receptors, BIBF 1120, also inhibits the parasite orthologs, leads to metacestode inactivation, and prevents parasite development of stem cell-containing primary cell cultures. This opens new ways for the development of anti-*Echinococcus* drugs using the parasite FGF receptors as target molecules.

## Acknowledgements

The authors thank Monika Bergmann and Dirk Radloff for technical support.

## Supporting information captions

**Fig S1. Amino acid sequences of *E. multilocularis* FGF receptors.** Deduced amino acid sequences of all three *E. multilocularis* FGF receptors are shown. After the name of each receptor the GenBank accession numbers of the cloned and sequenced cDNAs are shown as well as the gene designation according to the *E. multilocularis* genome project. Predicted signal peptides are marked in red, predicted IG-domains are marked in blue, tyrosine kinase domains are marked in yellow. Transmembrane regions are underlined.

**Fig S2. Amino acid sequence alignment of FGF receptor kinase domains.** Depicted is a CLUSTAL W amino acid sequence alignment of the tyrosine kinase domains of *E. multilocularis* EmFR1 (GenBank accession no. LT599044), EmFR2 (LT599045) and EmFR3 (LT599046) with those of *Schistosoma mansoni* FGFRA (SmFGFRA; Wormbase accession number: Smp_175590.1) and FGFRB (SmFGFRB; Smp_157300.1), *Dugesia japonica* FGFR1 (DjFGFR1; NCBI accession no.: Q8MY86) and FGFR2 (DjFGFR2; BAB92086.1), and the human FGFR1 receptor (HsFGFR1; NP 075598.2). Amino acid residues that are identical to the consensus of all sequences are printed in white on black background. Numbering to the right starts with amino acid 1 of the tyrosine kinase domain. The insert region of the split tyrosine kinase domain is indicated by a grey bar. Amino acids known to be highly conserved among tyrosine and serine/threonine kinases [34] are indicated by asterisks above the alignment. The two tyrosine residues known to be important for full activation of the human FGF receptor [35] are marked by black triangles. The TKD DFG motif which was modified to DNA to generate kinase-dead FGF receptors is indicated by black stars below the sequence.

**Fig S3. Expression of *E. multilocularis* FGF receptor genes in larval stages.** Indicated are fpkm (fragments per kilobase of transcript per million mapped reads) values for *emfr1* (red), *emfr2* (orange), and *emfr3* (blue) in *E. multilocularis* primary cell cultures after 2 days of incubation (PC2), after 11 days of culture (PC11), in mature metacestode vesicles without (MV-) and with (MV+) brood capsules as well as in dormant (PS-) and activated (PS+) protoscoleces. Transcriptome data have been generated during the *E. multilocularis* genome project and were mapped to the genome as described in [14].

**Fig S4. WMISH analysis of *emfr3* expression during metacestode development.** In all panels the WMISH signal is shown in green, and DAPI nuclear staining is shown in blue. **A.** Sense probe (negative control). **B.** Early brood capsule formation. **C.** Early protoscolex formation. **D.** Detail of early protoscolex formation, showing an *emfr3*+ cell in the region of the stalk connecting the developing protoscolex to the brood capsule. **E.** Detail of early protoscolex formation, showing the morphology of an *emfr3*+ cell (inset: only DAPI channel is shown). **F.** Late protoscolex formation. *bc*, brood capsule; *gl*, germinative layer: *ps*, protoscolex; *r*, rostellum; *s*, sucker; *st*, stalk. Bars: 25 µm for B, C, D and F; 10 µm for E.

**Fig S5. Phosphorylation of *E. multilocularis* FGF receptors upon expression in *Xenopus* oocytes.** *E. multilocularis* EmFR2 (EmFR2) and a kinase-dead version of EmFR2 (EmFR2 TK-) were expressed in *Xenopus* oocytes and stimulated with either 10 nM FGF1 or 10 nM FGF2 as indicated. After stimulation, cell lysates were generated, separated by SDS-PAGE and analysed by Western blot using antibodies against the myc-tag (Anti-myc; loading control) or phosphorylated tyrosine (Anti Ptyr). Depicted are the results for EmFR2. Results for EmFR1 and EmFR3 were comparable.

